# Single-cell heterogeneity in ribosome content and the consequences for the growth laws

**DOI:** 10.1101/2024.04.19.590370

**Authors:** Leandra Brettner, Kerry Geiler-Samerotte

## Abstract

Across species and environments, the ribosome content of cell populations correlates with population growth rate. The robustness and universality of this correlation have led to its classification as a “growth law.” This law has fueled theories about how evolution selects for microbial organisms that maximize their growth rate based on nutrient availability, and it has informed models about how individual cells regulate their growth rates and ribosomal content. However, due to methodological limitations, this growth law has rarely been studied at the level of individual cells. While *populations* of fast-growing cells tend to have more ribosomes than *populations* of slow-growing cells, it is unclear whether individual cells tightly regulate their ribosome content to match their environment. Here, we employ recent groundbreaking single-cell RNA sequencing techniques to study this growth law at the single-cell level in two different microbes, *S. cerevisiae* (a single-celled yeast and eukaryote) and *B. subtilis* (a bacterium and prokaryote). In both species, we observe significant variation in the ribosomal content of single cells that is not predictive of growth rate. Fast-growing populations include cells exhibiting transcriptional signatures of slow growth and stress, as do cells with the highest ribosome content we survey. Broadening our focus to non-ribosomal transcripts reveals subpopulations of cells in unique transcriptional states suggestive that they have evolved to do things other than maximize their rate of growth. Overall, these results indicate that single-cell ribosome levels are not finely tuned to match population growth rates or nutrient availability and cannot be predicted by a Gaussian process model that assumes measurements are sampled from a normal distribution centered on the population average. This work encourages the expansion of growth law and other models that predict how growth rates are regulated or how they evolve to consider single-cell heterogeneity. To this end, we provide extensive data and analysis of ribosomal and transcriptomic variation across thousands of single cells from multiple conditions, replicates, and species.

## INTRODUCTION

Growth laws describe robust correlations between cell growth and other phenotypes^1–4^. Perhaps one of the most well-studied growth laws pertains to the relationship between cell growth and the ribosome^5,6^. Numerous studies have shown that faster growing populations of cells possess more ribosomes^7–9^. This makes intuitive sense as ribosomes make most of the components that cells need to grow and replicate. However, ribosomes are expensive, being composed of many proteins and rRNA molecules, and comprising a large portion of the transcriptome^10–14^. Thus, cell populations are thought to tightly optimize ribosome levels to match their environment^15,16^. This tight optimization has been demonstrated at the transcriptional level; ribosomal as well as many other growth-supporting transcripts respond in concert to changes in nutrient abundance or environment quality^17,18^, sometimes within minutes^7^. The optimization of ribosome levels to match cell growth rate has been observed across diverse growth-limiting environments^19,20^ and organisms^7–9,21,22^, most often manifesting as a linear correlation^1,2,8^.

This tight coupling of population growth rate and ribosome levels has allowed important advances across diverse fields. It has enabled predictions of the growth of cells from the levels of their ribosomes or other growth-supporting molecules^7,9,23,24^, provided insights about the regulatory circuits that allow cells to fine-tune their growth to match their environments^16,19,25^, and suggested trends governing how cells across the tree of life should evolve to optimize ribosome utilization^1,2,26,27^. More generally speaking, the idea that cells calibrate their transcriptome and proteome to maximize their “cellular economy” has been a guiding principle across diverse biological disciplines^3,4,11^.

Although literature pertaining to the growth law discusses the ribosomal content per cell and models how single cells regulate their ribosome content^1,2,8,25^, the relationship between ribosomes and growth rate has most often been studied in aggregate. It has historically been difficult to measure the number of ribosomes in a single, tiny microbial cell. Alternatively, there are several well-established ways to macerate a large number of cells, measure the total amount of ribosome content in their lysate, typically as a mass-to-mass ratio of RNA to protein^6,8,28^, and how that quantity changes across similarly sized populations of cells growing at different rates. Numerous studies following this approach have shown that a population’s average ribosome level changes in lock step with its growth rate. But single-cell methodologies are revealing how taking the average can mask heterogeneity^29^. For example, this would miss rare subpopulations of cells that may have different growth strategies^30^, such as cells that prepare for environmental change by maintaining high levels of ribosomes despite growing slowly^31–33^, or by expressing stress-responsive proteins despite growing in optimal conditions^34^. Ignoring this heterogeneity may blind us to the existence of regulatory circuits that allocate resources in different ways in certain cell populations or environments. These circuits may represent different evolutionary solutions to the problem of maximizing fitness given resource availability. In sum, perhaps we are missing interesting biology by averaging away these signals.

Here we investigate heterogeneity in ribosome levels across single cells by utilizing data we and others recently generated by using a novel high-throughput method (Split Pool Ligation based Transcriptome sequencing, SPLiT-seq) that enables the sequencing of transcriptomes of individual microbial cells^35,36^. While more common cell isolation-based approaches have difficulty sequencing microbial cells due to their small size and unique anatomies, SPLiT-seq overcomes these limitations and allows the capture of thousands of RNA transcripts per microbial cell^35,36^. We implemented SPLiT-seq using random hexamer primers that recover rRNA, in addition to the typically used polydT primers that bias recovery towards mRNA^35,36^. This enables us to study how the ribosomal abundance of microbial cells changes as growth rate changes. We used data surveying roughly 8,200 haploid and diploid yeast cells (*Saccharomyces cerevisiae*) and 5,300 bacterial cells (*Bacillus subtilis*), sampled at different times – and thus varying growth rates – across the growth curve (**Figure 1a**). The cells averaged 6,390 or 2,644 rRNA reads per cell (for yeast and bacteria respectively).

**Figure 1.**
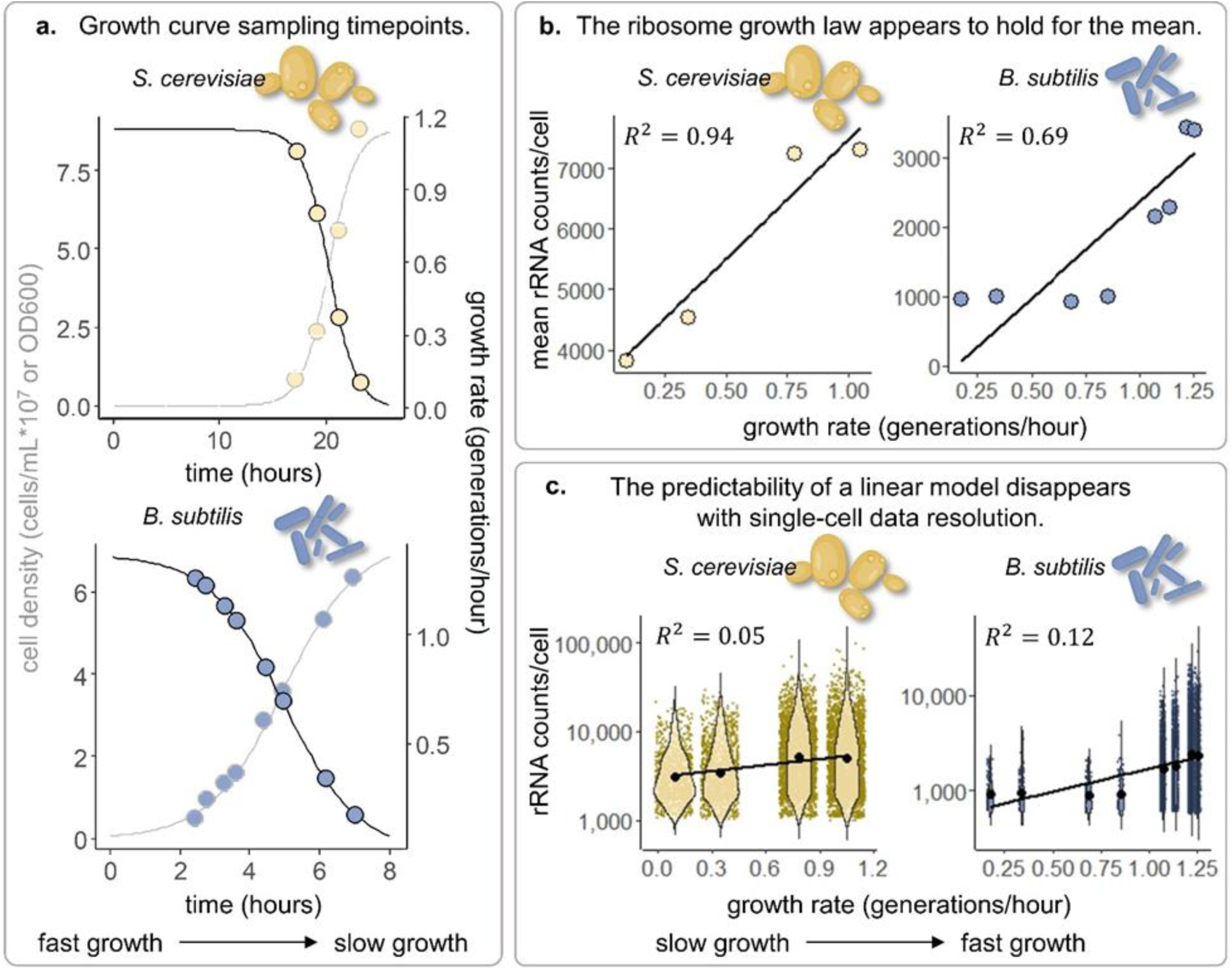
Population average ribosome levels correlate with population growth rates, but single-cell ribosome levels do not. **a.)** Yeast were sampled at four growth rates and bacteria at eight. Cell density (black) is measured in cells/mL*10^7^ in yeast and OD600 in bacteria. Growth rate is given in generations per hour (gray) against time in hours. Yeast were sampled at 0.92, 2.7, 5.7, and 8.6*10^7^ cells per milliliter. Bacteria were sampled at OD600 0.5, 1.0, 1.3, 1.6, 2.0, 3.6, 5.3, and 6.0. Yeast and bacteria cartoon icons were created using Biorender^93^. **b.)** Mean detected rRNA counts per cell correlate with the population growth rate in generations per hour. Trend lines and R^2^s were calculated by applying a linear regression model to the mean data points in R. **c.)** rRNA counts per cell do not correlate with population growth rate and are highly variable. rRNA counts for all cells are represented as violin distributions. Means (black dots) and trendlines are the same as in b. R^2^s were calculated by first applying a linear regression model to the entire dataset, and then confirming that the residuals were approximately normally distributed (absolute value Pearson median skewness < 0.3).

Contrary to the expectation that cells optimize their ribosome content to match their growth rate, we find enormous variation in single-cell ribosome levels across genetically identical cells that have the same access to growth-supporting nutrients. Like other studies^6,8,20–22,31^, we find that the average ribosomal content of a cell population is a strong predictor of the population growth rate. But the ribosomal content of a single cell cannot predict whether it was sampled from a fast or slow growing population. The variation in single-cell ribosome content we observe is inconsistent with what we would expect if it were driven by sampling noise, cell size, or single-cell growth rate. However, this variation is consistent with the findings of a recent time-lapse microscopy study in another bacterial species (*E. coli*) that finds single-cell growth rates and ribosome concentrations are uncorrelated^37^. Indeed, other growth laws, such as the correlation between growth rate and cell size, have also been shown to break down at the single-cell level^38^. In our study, we expand beyond growth rates to show that ribosome abundance is also not predictive of a single cell’s transcriptional state. For example, we observe cells with transcriptional signatures of stress and slow growth despite having high ribosome levels, and cells with low ribosome levels despite having transcriptional signatures of fast growth. Our work suggests caution when using a single cell’s ribosomal content as a predictor of its growth rate or behavior. This conclusion is relevant because measuring single-cell ribosomal content is becoming more common^33,39–41^, as are models that equate single-cell ribosome partitions with single-cell growth rates^42^. Our findings – demonstrating that variation in single-cell ribosome abundance is high, heavy-tailed, and exaggerated in fast-growing cell populations – can facilitate more accurate models of how cells regulate their ribosomal content.

In addition to our findings on variation in single-cell ribosome levels, we also observed another phenomenon that may be unexpected given a growth law where cells optimize their transcriptomes to maximize their growth rates^3,15,43^. We observed transcriptional heterogeneity in populations of genetically identical cells under nutrient rich conditions. For example, even when we sample cells very early in their growth curve, we observed some bacterial cells expressing transcripts related to sporulation, some yeast cells inundated with retrotransposons, and some cells of both species expressing canonical stress responses. These observations may not be surprising to some scientists because cell-to-cell variation seems to be the rule rather than the exception^44–48^. But on the other hand, heterogeneity is somewhat inconsistent with the idea that microbial cells employ a single strategy – optimizing their phenotype for rapid growth – to maximize fitness in nutrient rich conditions. More and more work is revealing alternate evolutionary strategies including dormancy^30^ and bet hedging^34,49–51^ to the extent that more inclusive definitions of evolutionary fitness are emerging^52^. This work, and ours, highlights how further modeling efforts are likely needed to reconcile ideas about optimization of the cellular economy to support fast growth with observations of phenotypic heterogeneity.

## RESULTS

### Single-cell ribosome levels do not correlate with population growth rate

Either yeast (*S. cerevisiae*)^35^ or bacteria (*B. subtilis*)^36^ were previously grown in batch culture and sampled over time starting when populations were at or near steady state growth and ending as the cells were exhausting their nutrients (4 samples over time for yeast, 8 for bacteria). For each sample, cell counts or optical density were measured^35,36^. From these measurements, we created growth curves for each population of cells (**Figure 1a**, gray lines). To track the rate at which the populations of cells were growing at each time point, we calculated a sigmoidal function to fit the growth curve and then took the log2 ratio between the adjacent data points (**Figure 1a**, black lines). We used these calculations to estimate the instantaneous growth rate of each population we study in doublings or generations per hour (**Figure 1b-c**, horizontal axes). These cells sampled at each time point were also used to perform single-cell RNA sequencing via Split-Pool Ligation based transcriptomics sequencing (SPLiT-seq)^35,36^. We used these transcriptional data to calculate the ribosomal content of, on average, 2,045 and 683 single cells per sample for yeast and bacteria, respectively (**Figure 1b**, vertical axes). As expected according to the growth law, there is a strong correlation between the average amount of ribosomal RNA in a population of cells and that population’s instantaneous growth rate (**Figure 1b**). In both yeast and bacteria, we find that population growth rate explains 94% and 69% of the variance in average rRNA abundance respectively when using a linear model (**Figure 1b**, R-squareds). The non-linearity in slow-growing *B. subtilis* populations, in which ribosomal content appears to be less responsive to changes in nutrient content, has been observed for other bacterial species as well^22^.

However, this correlation breaks down at the single-cell level where the population growth rate explains 5% and 12% of the variance in single-cell rRNA abundance in yeast and bacteria respectively (**Figure 1c**, R-squareds). Even in conditions when cells are flush with nutrients and the population is doubling at the fastest rate we survey, there is massive variation in ribosome content across cells spanning more than two orders of magnitude (**Figure 1c**). Within a single timepoint, there is up to 25 times as much variation as we observe on average across timepoints (**Figure 1b**). The high level of variation we observe seems unexpected if individual cells optimize their ribosome levels to maximize their growth rate given the nutrient availability. This variation is inconsistent with that which we expect if it were driven by sampling noise, cell cycle state, or cell size (**Figure S1 – S3**). For example, our analysis includes both haploid and diploid strains of yeast, which are known to differ in size by approximately 1.5X on average^53^. And yet, ploidy explains less than 1% of the variance in single-cell rRNA abundance in a categorical linear model (**Figure S2**). Moreover, our further analyses suggest that ribosome content is a poor predictor of single-cell growth rate and/or transcriptional state (**Figures 2 – 5**).

**Figure 2.**
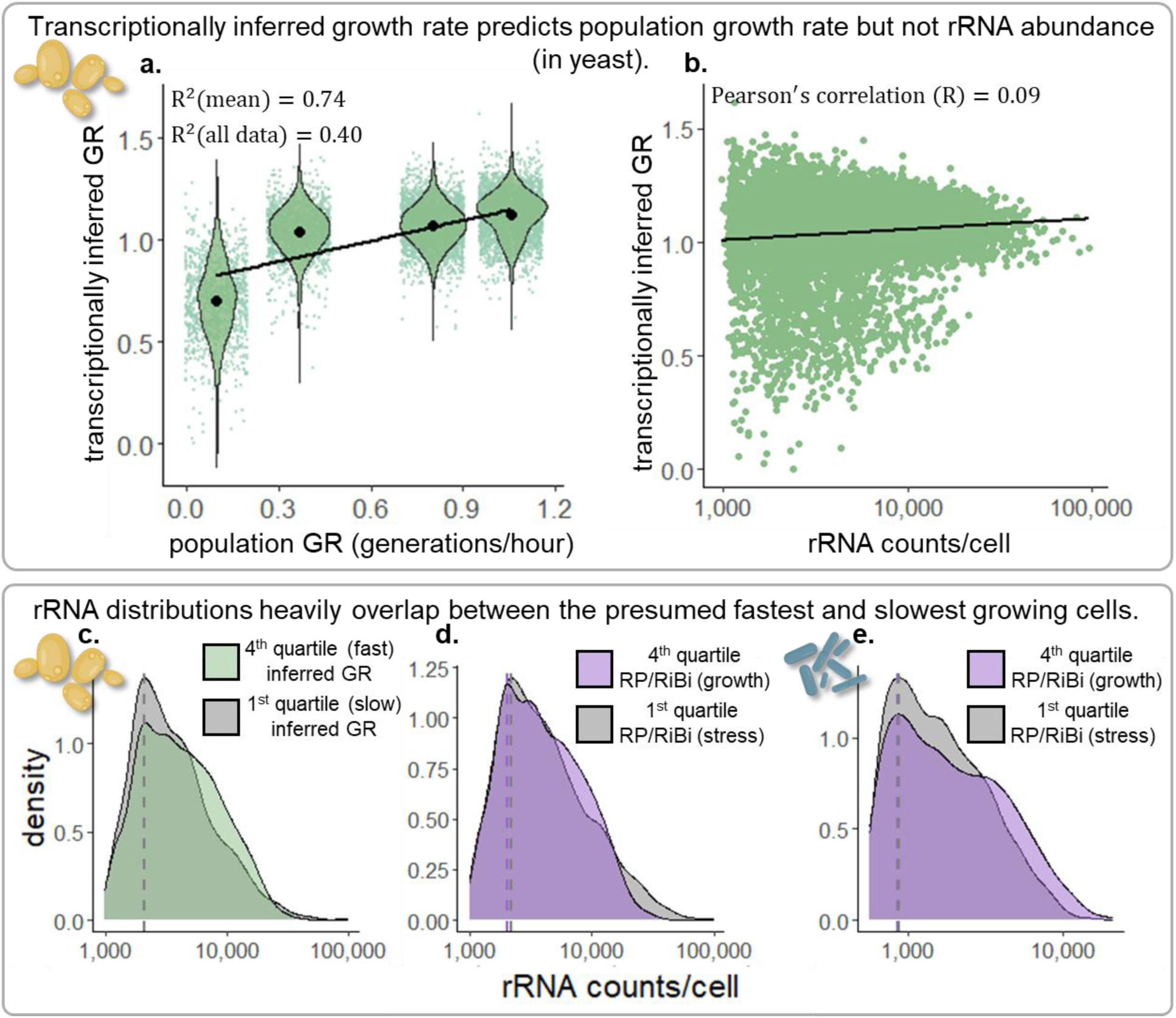
rRNA abundance in cells with growth associated transcriptional signatures does not differ from cells with stress signatures. **a.)** Means and single-cell distributions of transcriptionally inferred growth rates against population growth rates (in yeast). R^2^s were calculated by applying a linear regression model to the means (black dots) and all data for each single cell (green dots). **b.)** Transcriptionally inferred growth rate for each cell does not strongly correlate with corresponding rRNA counts. **c.)** rRNA counts/cell density distributions for the cells with the 25% highest (green) and lowest (gray) inferred growth rates (in yeast) are largely overlapping. Dotted lines represent the density distribution maximum, or the largest mode. **d.) and e.)** Similarly overlapping distributions are observed for the cells with the highest percentage of the transcriptome dedicated to ribosomal proteins and ribosomal biogenesis genes (lavender) versus stress genes (gray) as defined in yeast^18^, and a similar cohort of genes for *B. subtilis* (Supplemental Table 1).

### Single-cell ribosome levels do not correlate with transcriptionally inferred single-cell growth rates or expression states

Because we sample from the entire transcriptome, including mRNA as well as rRNA, we can ask questions about how the expression profiles of pertinent mRNA transcripts correlate with rRNA abundance. Despite surveying numerous modules of gene expression that are indicative of growth rate and/or stress responses, we were unable to find the expected differences in the rRNA content of cells in different transcriptional states (**Figure 2**). In other words, cells with mRNA signatures indicative of fast growth are indistinguishable from cells with mRNA signatures indicative of slow growth or stress in respect to their rRNA content (**Figure 2**).

Previous work in yeast has defined core sets of genes that respond to changes in growth rate^9,23,54^. The expression levels of these genes are so correlated with growth that they can be used to predict the growth rate of yeast populations in novel conditions, including steady-state conditions where growth is limited by different nutrients, and batch culturing conditions where growth rate declines as nutrients expire^7,9,23,54^. We find that these genes are also predictive of population growth rates in our yeast populations (**Figure 2a**). However, the levels of these transcripts are not strongly correlated with rRNA abundance (**Figure 2b**). The lack of correlation between a cell’s detected rRNA and the genes it was transcribing at the time we sampled it is even more apparent when we compare the top quartile (25%) of cells with the fastest transcriptionally inferred growth rate^9^ (**Figure 2c**, green) to the quartile of cells with the slowest inferred growth rate (**Figure 2c**, gray). The distributions of ribosome content for these two populations are entirely overlapping with almost identical maximum densities (**Figure 2c**). We find the same results when we use other (overlapping) sets of genes to define yeast cells with signatures of fast growth (ribosomal proteins and supporting genes) versus slow growth (stress-responsive genes)^18^ (**Figure 2d**), and when we perform a similar analysis in bacteria (**Figure 2e**).

In sum, the number of ribosomal RNAs within a cell (**Figure 2**) does not seem to report useful information about the cell’s transcriptional program or inferred rate of growth. This finding is consistent with recent work using time-lapse microscopy that also demonstrates no correlation between a cell’s ribosomal protein levels and its rate of growth^37^. These findings challenge the conventional assumptions about the relationship between ribosomal RNA content and growth rates. Importantly, these observations warn against measuring single-cell ribosome levels to infer single-cell growth rates or population growth dynamics^42^, which is becoming increasingly attractive as methods for single-cell quantification improve in precision and throughput^33,39–41^.

### Variation in single-cell ribosome levels is highest when variation in single-cell growth rates is lowest

If single-cell rRNA levels determine single-cell growth rates, then we would expect variation in growth rates is highest when variation in rRNA levels is also at its peak. But that is not what we see (**Figure 3**). By using two separate approaches, we show that variation in single-cell growth rate is negatively correlated with population growth rate (**Figure 3a, b**). We initially investigated variation in single-cell growth rates by inferring these growth rates from gene expression profiles in yeast, as we did in **Figure 2**. We also obtained an independent data set of single-cell *S. cerevisiae* direct growth measurements where thousands of yeast microcolonies were monitored over time using high-throughput microscopy^55^. We aggregated the growth measurements and calculated the variation using the coefficient of variation (mean normalized standard deviation, σ/µ) of the individual growth rates of the single cells and microcolonies as stratified by time point. For both datasets, both inferred and directly measured, the variation of single-cell growth rates declines as the population average increases (**Figure 3a, b**).

**Figure 3.**
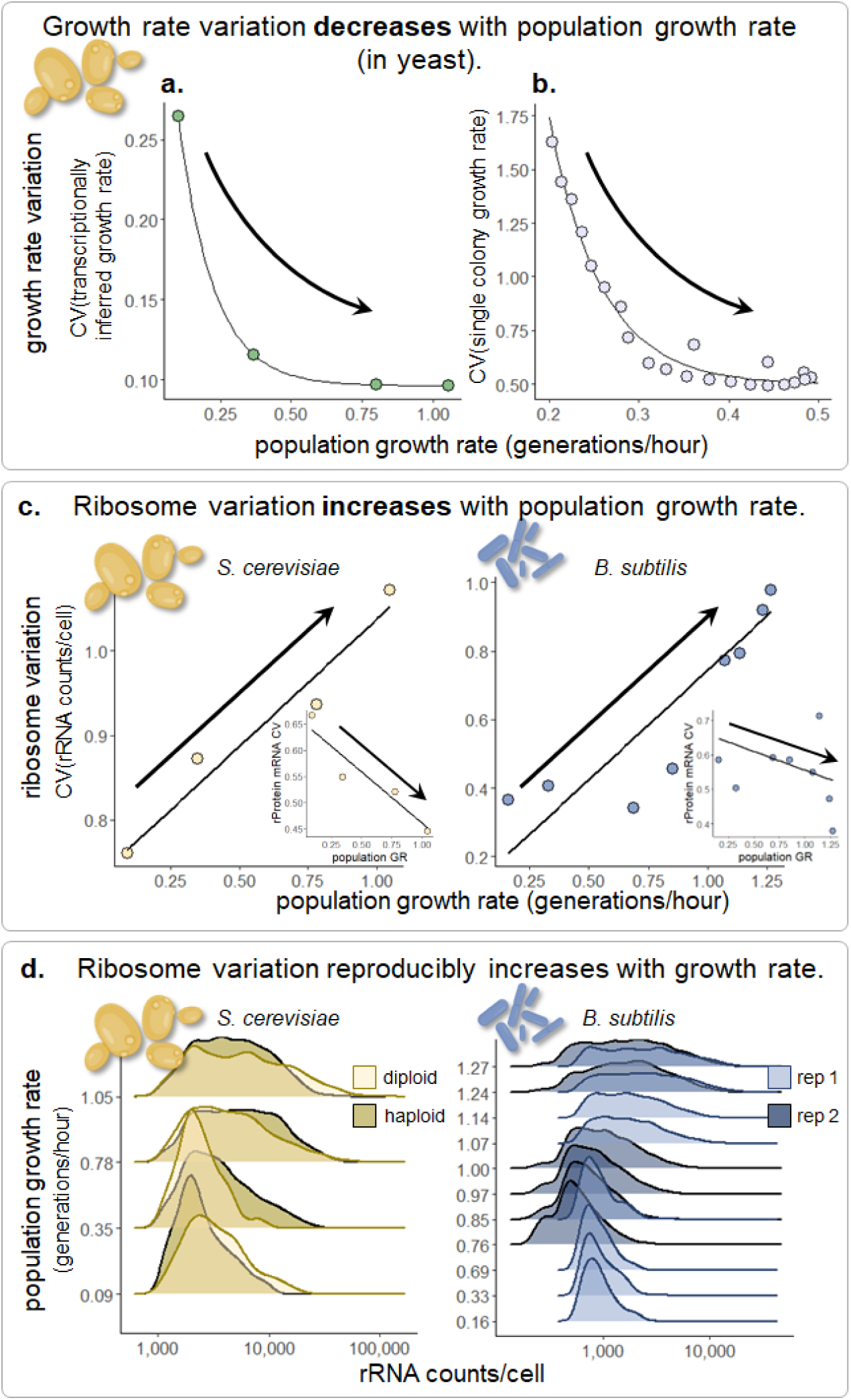
Variation in single-cell growth rates does not explain the observed variation in rRNA abundance. **a.)** Coefficient of variation (σ/µ) of single-cell growth rates estimated using gene expression (Figure 2a-c) for cells in each timepoint decreases with population growth rate. **b.)** Coefficient of variation of single microcolony growth rates decreases with the average growth rate for each imaging time^55^. **c.)** Coefficient of variation of ribosome counts for cells in each timepoint increases with population growth rate. Inset plots show the CV for ribosomal protein mRNA counts/cell. Trendlines are shown to indicate the direction of correlation with population growth rate. **d.)** rRNA counts/cell densities for each timepoint split by ploidy in yeast and biological replicate in bacteria.

Conversely, we observe that variation in rRNA abundance correlates positively to the population growth rate. That is to say, fast growing populations of cells, sampled when nutrients are most abundant, have the highest variation in rRNA counts in both yeast and bacteria (**Figure 3c**). This positive correlation appears to be consistent with at least one previous study in another organism^22^. This trend also persisted for another measure of data variance or spread that is not reliant on normal data distributions (**Figure S4**). However, this trend does not persist when we analyze other growth-supportive molecules such as ribosomal protein mRNAs (**Figure 3c**; insets). This observation may suggest that there is something unique about the biology of ribosomal RNA that causes variation in its levels to be highest when the population growth rate is also at its highest.

In sum, our analyses suggest that single-cell growth rates are most variable in slow-growing populations of cells (**Figure 3a, b**), but single-cell ribosome levels are most variable in fast-growing populations of cells (**Figure 3c**). These results run counter to the hypothesis that single-cell growth rate variation underlies the variation we observe in ribosome abundance. This result provides additional support for our conclusion that single-cell rRNA levels are not correlated with single-cell growth rates.

These results, specifically, the pattern of widening rRNA distributions with population growth rate, are reproducible across both haploid and diploid yeast, and two biological replicates in bacteria (**Figure 3d**). We also observe that, contrary to the typical assumption of biological systems, the variation in single-cell ribosome content is not normally distributed and has a very long tail (**Figure 3d**, right tail is even longer than it appears because these plots are in log scale). Previous work has shown biological models that expect variation to be normally distributed around the mean can fail to describe the behavior of single cells^56^. Our findings suggest a complex relationship where ribosomal RNA content varies independently of growth rates, highlighting the need for refined biological models to better understand single-cell dynamics. We hope that by measuring the shape of these single-cell distributions (**Figure 2a & 3d**), our work will enable more precise models of the biology underlying how single-cells regulate their ribosome content, growth rates, and other behaviors.

### Cells with the most ribosomes do not all have transcriptional signatures of faster growth, and cells with the fewest ribosomes do not all have signatures of slower growth

As a final demonstration that the ribosomal content of a cell is not predictive of its rate of growth or transcriptional state, we decided to compare “apples to apples,” so to speak. We ordered the cells by their ribosome abundance from least to greatest across all time points. If ribosomes are the dominant signatures of growth rate, we would expect the cells with the most ribosomes to uniformly exhibit growth signals in their transcriptomes, and the cells with the least ribosome abundance to show signs of stress or slow growth.

We separated the cells into four quartiles and used Louvain clustering to investigate any subpopulations that might be present within each quartile. Differential expression analysis on the identified subpopulations revealed that, within each quartile, cells segregated by signatures of growth and stress (**Figure 4a**). Contrary to our expectations given the growth law, the cells with the highest number of rRNAs do not all have transcriptomes suggestive of fast growth (**Figure 4a, i-ii**). Similarly, some cells with the fewest ribosomes still appear to be employing transcriptional programs associated with a growth-oriented directive (**Figure 4a, iii-iv**).

**Figure 4.**
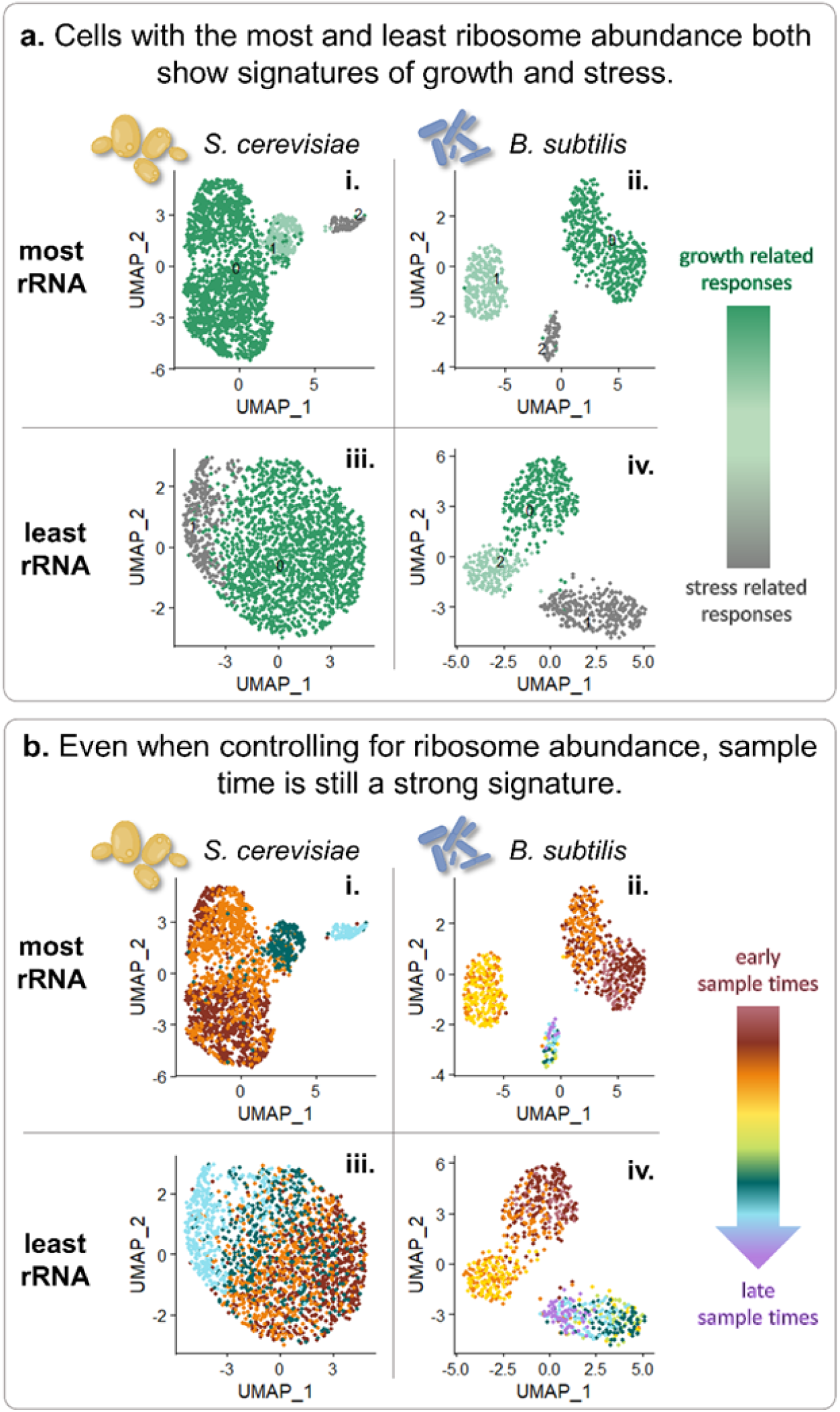
Cells show differential subpopulations and signatures of sample timepoint despite having similar ribosome levels. **a.)** UMAP embeddings of cells from the most (i-ii) and least (iii-iv) rRNA abundance quartiles colored by the Louvain cluster contain both cells with transcriptional signatures of growth as well as stress. **b.)** UMAP embeddings of cells from the most (i-ii) and least (iii-iv) rRNA abundance colored by sample timepoint suggest that having similar amounts of rRNA does not necessarily make cells similar at the rest of the transcriptional level.

To be clear, in this analysis, we did not *a priori* define growth or stress responses as we did previously in **Figure 2**. Instead, we used an algorithm to cluster cells with similar transcriptomes, and then subsequently asked what transcriptional differences defined each cluster (**Figure 4a**). In yeast, the clusters which we labeled “growth” (**Figure 4a**, green) are defined by the expression of genes involved in glycolysis, ribosome biogenesis, and translation. The clusters which we labeled “stress” (**Figure 4a**, gray) show upregulated gene expression of heat stress, osmotic stress, and protein misfolding stress pathways. In bacteria, in the “growth” clusters (**Figure 4a**, green) we see differential expression of similar translation and ribosome biogenesis pathways, and also operons involved in the import and digestion of preferred carbon sources. The “stress” cluster (**Figure 4a**, gray) is defined by expression of genes such as sporulation initiation, bacteriocin production, and surfactin production. Additionally, in both organisms we see an intermediate cluster of cells (**Figure 4a**, light green) that are shifting from glycolysis and preferred carbon digestion to the TCA cycle and operons for the import and digestion of less desirable carbon sources such as short chain fatty acids, and in *B. subtilis*, the lichenan pathway^57^. The salient observation is that all clusters (stress and growth) are present in all quartiles, including those containing cells with the most ribosomes (**Figure 4a,** i-ii) and cells with the least ribosomes (Figure 4a, iii-iv). This finding challenges the assumption that ribosomal content is a reliable indicator of cell growth rate or state.

An alternate interpretation of this growth law might posit that cells with the highest or the lowest ribosome content include those with non-optimal numbers of ribosomes given their environment. In this case, one might expect cells that have an intermediate ribosomal content to express growth-responsive, rather than stress-responsive transcripts. However, this interpretation is inconsistent with our observation that both cell types (those expressing growth-responsive or stress-responsive transcripts) are present in quartiles of cells possessing intermediate amounts of ribosomes as well (**Figure S5**). Further even after we sort by ribosome abundance, the strongest signature in the clustering is the sample time point (**Figure 4b**). This suggests that a cell in a quartile of ribosomes from early in the growth curve and a cell with comparable ribosomes from later on are not most essentially defined by their ribosome abundance. Perhaps this result appears obvious in retrospect, but if ribosomes and growth activities were intrinsically linked as the growth law suggests, then the null assumption would be that these cells should be alike. Seemingly, having similar amounts of rRNA does not make cells similar at the rest of the transcriptional level. Overall, our findings advise caution when equating an individual cell’s ribosome level to its growth rate, or more generally, to any particular cell state or behavior.

### Is the challenge rooted in the metric or the growth law model?

Our findings reveal that single-cell rRNA abundance does not well predict single-cell growth rate (**Figures 1 - 4**), echoing recent work in *E. coli* that shows no correlation between ribosomal protein abundance and growth rate^37^. This raises two possibilities. First, growth law models may be focusing on the wrong metric. The sheer number of ribosomes might be less indicative of growth than other cellular factors, such as the levels of mRNA transcripts that support growth^9^ or the abundance of active ribosomes^58^. Though many studies that validate the growth law in populations of cells, including this work (**Figure 1a**), measure ribosome content^8,28,31^, newer research suggests that mRNA concentration— an indicator of ribosome activity—may offer an alternative model for cell growth^59^. We modeled this as the mRNA/rRNA ratio in single cells, and we see a stronger, but still not predictive correlation with growth rate or cell state (**Figure S6**). Curiously, we find differences in mRNA/rRNA ratio between replicates that are supposed to be the same. Specifically, we see that one replicate of our bacterial experiment harbors a subpopulation of cells with very high mRNA/rRNA ratios and upregulation of sporulation genes (**Figures 5, S7, S8**). This difference between replicates, and further, the presence of a sporulating cell subpopulation within a replicate, caused us to wonder whether the core issue with the growth law lies not in the metric, but in the model’s assumptions. Current growth law frameworks typically assume uniformity in cellular behavior across populations, positing that cells are fine-tuning their transcription to reach optimal levels of growth-related molecules^3,15,43^. But a growing body of single-cell work suggests that populations of cells are rarely uniform in their phenotypes^36,45,46,60^. This prompted us to further investigate whether there is evidence of heterogeneous cell states in our study, even among cells subjected to identical nutrient conditions.

### The transcriptional heterogeneity we observe is inconsistent with a model in which all cells are optimizing their cellular economy for a growth-oriented objective

While the above analyses suggest that there is a great deal of variation in rRNA among cells exposed to the same environment and nutrient levels, a remaining question is if such cells also exhibit transcriptionally distinct states. Heterogeneity of this nature seems potentially inconsistent with the spirit of the growth law, which is based on the idea that cells have evolved to conform to a singular evolutionary strategy: tightly optimizing their cellular economy for maximal growth^3,4,11^. In multiple sampled timepoints, and in both yeast and bacteria, we see what appear to be transcriptionally distinct cell types co-existing in the same sample (**Figure 5a).**

This heterogeneity appears to be meaningful. In both yeast and bacteria, we observe cells that show the expression signatures associated with faster growth, such as upregulation of genes related to ribosome production, translation, carbon utilization, RNA processing, and cell replication (**Figure 5a**; green). However, we also find significant numbers of cells from the same environments that show signs of classic stress responses, such as upregulation of general stress proteins, as well as genes related to respiration, the TCA cycle, mitochondrial proteins, trehalose metabolism^34^, and low glucose conditions (**Figure 5a**; gray). In addition to these co-existing subpopulations that are defined by transcriptional signature of growth versus stress, we also see evidence of additional coinciding transcriptional states. For example, in some samples we see subsets of yeast cells that express a high abundance of retrotransposons, comprising up to 42% of their transcriptome (**Figure 5b**). Similarly, some bacterial samples contain clusters of cells that appear to be sporulating even when nutrients are abundant (**Figure 5c**). Cells with these phenotypes are not optimized for growth, either because they are diverting exorbitant resources towards producing a genetic parasite (retrotransposons)^61^ or because they have entered a costly and ‘irreversible’ process of non-growth (sporulation)^62^. And yet cells expressing stress- and spore-associated transcripts or retrotransposons are present in every time point we survey, including when nutrients are most abundant, and cells have just exited steady-state growth (**Figures 5, S8**). These cells represent the antithesis of a growth-oriented objective, calling into question the applicability of a growth law model that optimizes all cell’s transcriptomes to support the maximum rate of growth.

**Figure 5.**
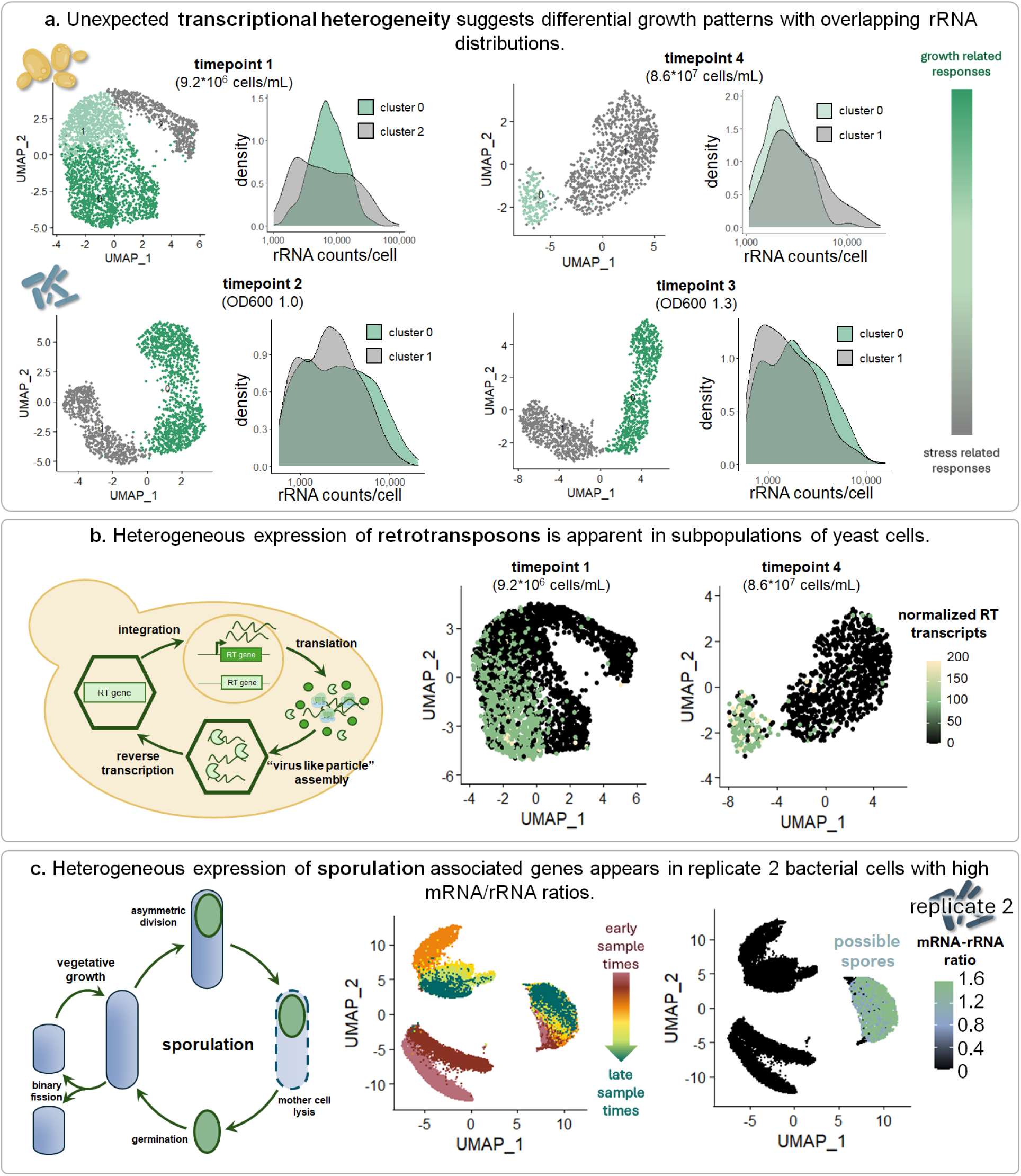
Unexpected subpopulations are observed across the growth curve in both yeast and bacteria. **a.)** UMAP embeddings of cells from the first timepoint and the last timepoint of the yeast growth curve and timepoints 2 and 3 in bacteria, colored by Louvain cluster. Green clusters include cells with differentially expressed genes more associated with growth and gray clusters with differentially expressed genes associated with stress. Density plots show that the distributions of rRNA counts/cell for green and gray clusters overlap. **b.)** UMAP embeddings of cells from the first timepoint and the last timepoint of the yeast growth curve colored by retrotransposon gene transcript abundance. Retrotransposons are mobile genetic elements within the yeast genome typically thought to be activated by stress^94^. However, here we observe cells expressing them not only in the final timepoint of the yeast growth curve, but also in the first timepoint in cells with transcriptional signatures of fast growth. **c.)** UMAP embeddings of bacterial cells from an independent replicate experiment^36^ colored by sample time (left) or mRNA-rRNA ratio (right). Cells in the UMAP visual cluster with distinctly high mRNA-rRNA ratios also upregulate genes related to sporulation, and are present in all timepoints, including when cells are growing in rich media. Cartoon diagrams inspired by^61,95^.

While our observations beg questions about why this heterogeneity exists (**Figure 5**), our goal here is simply to point out that it does. And that its presence limits the utility of summary statistics such as the population average. When meaningful heterogeneity is present, biological models based on population averages often fail to make accurate predictions about the behavior of individual cells^29,56^. The heterogeneity we observe is consistent with other studies that show single-cell heterogeneity is the rule rather than the exception^34,36,44,45,55,63^. Though high-throughput single-cell transcriptomics in microbial cells is just emerging, previous work examining the transcriptomes of a small number of cells or higher-throughput studies using fluorescent markers have shown that heterogeneity is incredibly common. For example, previous work suggests that persister cells^47,64^, or more generally, slow growing cells that have unusually high stress tolerance^34^, exist even in fast-growing cell populations that have access to high levels of nutrients. Even under conditions of stress, when mounting a transcriptional stress response may mean the difference between life or death, not all cells respond uniformly, or even respond at all^46,56^.

Single-cell heterogeneity is sometimes thought to reflect bet-hedging, or a simultaneous display of multiple evolutionary strategies to prepare for an unknown future^34,51,65^. For example, an alternate evolutionary strategy besides maximizing growth rate may involve harboring excess ribosomes to prepare for non-growth conducive conditions that require increased translation^28,31,66,67^. Another strategy may involve expressing stress-responsive, rather than growth-responsive transcripts, to prepare for unexpected and unfavorable environments^34,47,64,65^. But cell-to-cell heterogeneity could also simply result from unavoidable processes like expression stochasticity, DNA damage, or aging^45,60,68^. Whatever the underlying cause of the heterogeneity, our observation that not all single cells optimize their ribosome content (or their transcriptomes) in the same way to match their nutrient level is significant. The heterogeneity we observe within samples, as well as the differences we and others^69–71^ observe between replicate experiments, highlight lingering mysteries surrounding how cells regulate their phenotypes in response to environmental stimuli. We wonder to what extent we can begin to shed light on these mysteries by expanding growth law models beyond population averages.

## DISCUSSION

We utilized a novel single-cell method^35,36^ to study the ribosome growth law, or the robust correlation between collective ribosome content and population growth rate. Similar to previous studies^6,8,28,31^, we observed this relationship holds true in that the average ribosome level of a population correlates with its growth rate (**Figure 1b**). However, our data, along with recent studies, fail to recapitulate this law^37^, or other growth laws^38^, at the single-cell level (**Figure 1c**). This raises the general question: how can a phenomenon that is apparent in aggregate be absent across individuals? This scenario, while counterintuitive, is not necessarily uncommon; one classic example is Simpson’s paradox^72^.

### How can the average fail to predict the behavior of individuals?

Consider, for instance, a toy example where at all times there exist two subpopulations of cells each with different ribosome contents. The correlation between a population’s average growth rate and ribosome content could then reflect changes in the frequency of each subpopulation, rather than a continuous scaling of growth with ribosome content at the single cell level. This bimodality in ribosomes has been observed during starvation in other organisms^33^, but we do not find obvious evidence for this in our study. Even when we classify cells into different transcriptional subpopulations, we cannot find a clear difference in their ribosome content (**Fig 2c-e & Figure 5a**, inset).

However, many other types of non-normally distributed data, in addition to the bimodal distribution described in this toy example, can result in scenarios where summary statistics such as the mean fail to describe the behavior of individuals^73^. Models based on population averages often result in poor predictions because they assume that variability among cells follows a normal distribution. This assumption can lead to inaccuracies, especially when the distributions of single-cell properties are asymmetric or heavy-tailed, as they are in our findings (**Figures 1c, 2c-e, 3d**). This means that, even if many cells are following a growth-oriented objective and striving to achieve an optimum concentration of ribosomes for maximal growth, that concentration might not be reflected in the average. A model based on the average might fail to capture regulatory nuances that cause tailed distributions^56^, such as those resulting from transcriptional bursting^74^. Fortunately, improved predictions of single-cell behavior can emerge upon constraining gene regulatory models using the full distribution of single-cell phenotypes rather than relying on summary statistics^56^. Given we measured these distributions, in replicate, for both yeast and bacteria in multiple conditions, perhaps there is potential to further develop more complex growth law models that describe how single cells regulate their ribosomal content, growth rate, and transcriptional state.

### What could we learn about growth regulation and evolution by considering complexity?

Perhaps even a stochastic model that can capture more accurate distributions of single-cell ribosomes is insufficient to make any substantive predictions about natural systems if it does not take a more holistic account of cellular phenotypes, realistic environments, and evolutionary histories. Reductionism, or the principle that we can understand complex problems by studying them in isolated, simplified conditions, has facilitated the unraveling of many knotty scientific questions. Yet, this approach can fail when applied to living organisms. Studying a fundamental, constant force like gravity in a vacuum enables you to eliminate confounding variables like air resistance. However, unlike gravity, biological systems are dynamic and mutable. When we place cells in a vacuum, so to speak, by exposing them to simplified laboratory environments, and especially continuous culture, we impose alternate selection pressures. In asking cells to grow as fast as they are able in a given nutrient, often one of the first things they do is irreparably break their regulatory systems, such as glucose sensing^75–77^, in the pursuit of rapid growth. These experiments, then, may be particularly susceptible to the “observer effect,” wherein the simple act of growing the cells in the lab fundamentally changes what we are trying to study. By irreversibly altering their behavior, any insights we gain may not be predictive of how cells function in nature.

In nature, cells exhibit heterogeneity in cell behavior, the ability to switch strategies, and adaptability to changing environments. The traditional growth law model postulates that cells are evolutionarily optimized to grow as efficiently as possible, particularly by tightly regulating ribosomal content or activity^8,16,25^. But it is not clear to what extent pressure to maximize growth rate is a dominant pressure in natural environments. Microbial life, in its tenure on Earth, has often existed in genetically diverse, fluctuating environments where maximal growth is not the sole priority, nor is selection always the dominant force for change. Population size and structure can change the influence of other driving forces such as drift, mutation, and gene flow^78–80^. Nature does not always produce the “best” or most efficient solution. Instead, life evolves under a balance of competing pressures, creating organisms that are the product of compromises and trade-offs. Our and others^8,28,31^ observation that the ribosomal content of a cell population correlates with its rate of growth suggests that pressure to utilize resources efficiently and maximize growth rate may be one of these competing pressures. But our and others^37,38^ observation that this correlation breaks down at the single cell level suggests a need to move beyond models that focus solely on growth maximization in order to understand not only how cells regulate their growth and transcriptional phenotypes, but also the pressures under which cells evolved.

## METHODS

### Growth curve and single-cell sequencing data

Cell culture methods were described in previously published works by L. Brettner and others^35,36^. Briefly, overnight liquid cultures from colonies of diploid s288c cells and haploid BY4741 *Saccharomyces cerevisiae* and PY79 *Bacillus subtilis* cells were started in cultures of ∼300 cells/mL in synthetic complete plus 2% glucose media for yeast or OD600 0.02 in Lennox broth media for bacteria respectively. These cultures were then sampled and measured systematically until the populations started to enter stationary phase. The samples were saved and processed for single-cell RNA sequencing using the SPLiT-seq method^35,36,81^. Sequencing data for these experiments have also already been published^35,36^. However, the saved *S. cerevisiae* sample libraries were re-sequenced on the NovaSeqX platform with paired end reads for a total of 300 cycles. This provided further and deeper coverage than what was reported in Brettner et al. 2024^35^. Sequencing and processed data for *S. cerevisiae* can be found through GEO at accession number GSE251966 and for *B. subtilis* at GSE151940. Additional processed data and analysis scripts can be found at the Open Science Framework: https://osf.io/kjfbz/. Additionally, there were two repeat experiments reported in Kuchina and Brettner et al. 2021^36^, described as M14 (replicate 2) and M15 (replicate 1). We focus most of our analyses on the data from M15 (replicate 1) as the data in M14 suggest many of the cells may be transitioning to a non-growing sporulation state (**Figure 5c and S8**).

### Cell density curve and growth rate curve calculations

The cell densities reported in cells/mL for yeast and OD600 for bacteria, and timepoints in hours were used to calculate a sigmoidal curve through a nonlinear weighted least-squares estimate in R (nls function). The model was fitted to the function for a sigmoid as follows (**Figure 1a**, gray lines).

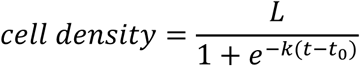

Where L is the saturation density, t0 is the half maximal time, and k is the slope. The data points from these models were then used to fit the growth rate curve in generations per hour using a log2 ratio between adjacent points normalized to the time increment as follows (**Figure 1A**, black lines).

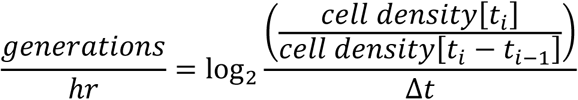

A sigmoidal function might not be the best approximation of growth under very low densities and early in the growth curve as it assumes continuous exponential growth and cannot incorporate any kind of lag phase. This might explain why the maximal estimated yeast growth rate is slightly higher than typically reported in the literature (∼1.05 generations/hour compared to ∼0.75 generations/hour). However, the absolute value is of little importance, and instead the shape of the curve and the relative relationships between the points are what impact the relevant downstream models (eg. the goodness of linear fits, R^2^).

### Single-cell RNA seq data processing, rRNA abundance calculations, and mRNA normalization

Sequencing reads were aligned to the R64 s288c genome for *Saccharomyces cerevisiae* from NCBI or the ASM904v1.45 genome for *Bacillus subtilis* from EnsemblBacteria and sorted by barcode using STARsolo^82^. As both the yeast and bacterial genomes have significant sequence homology, especially between rRNA genes, we enabled the uniform multimapping algorithm within STARsolo to keep overlapping reads. This outputted gene-by-barcode matrices representing the detected gene counts for every cell. To remove empty and low gene detection barcodes, we applied the “knee” detection filter previously described in Brettner et al. 2024^35^. These quality-thresholded gene-by-barcode matrices were then converted to the R datatype, Seurat Objects, using the Seurat R package for further analyses^83^.

rRNA genes were identified using the Saccharomyces Genome Database (SGD)^84^ and SubtiWiki^85^. Counts from these genes were then aggregated to calculate an rRNA abundance for each cell. While single-cell RNA sequencing data is typically highly sparse for mRNA, the minimum combined rRNA counts detected in one cell in these data were well above 500 reads. As scRNAseq data normalization methods are designed for sparse, low count data, we used the raw total rRNA counts in all ribosome abundance analyses to avoid unnecessary data distortion^86^, though the normalized and scaled rRNA data can been seen in Supplemental Figure 1

Normalization, scaling, nearest neighbor calculations, clustering, and differential expression analyses were performed similar to the tutorial provided by the Satija Lab^87^ on mRNA datasets with the rRNA removed. As rRNA typically makes up greater than 90% of the detected RNA reads and clustering is relative, keeping it in made for skewed clustering analyses that obscured some of the interesting biological behaviors.

### Ruling out sampling noise, cell size variation and cell cycle variation as explanations for the observed variation in rRNA counts/cell

While many factors likely contribute to variation in ribosome levels, we could not find any artifact that could easily explain away most of this variation. For example, the variation we see is not consistent with sampling noise. If most of the variation is resultant from sampling noise, then we would expect relative noise to be consistent or to be highest in slow-growing populations of cells that have on average, fewer ribosomal transcripts. However, that is not the case (**Figures 1 & 2**). Relatedly, we showed that normalized measures of single-cell ribosomal content, besides absolute rRNA abundance (**Figure 1**), also do not predict the population’s growth rate and nutrient access (**Figure S1**). For example, normalizing each cell’s mRNA content by its rRNA content may reduce sampling noise while also serving as a general estimate for the portion of each cell’s ribosomes that are active, which recent work shows also correlates with growth rate^58,59^. mRNA is a highly transient molecule in microbes and should be readily occupied by ribosomes^88,89^. If the mRNA/rRNA ratio is high, we expect that the cell is metabolically active, since it is dedicating more of its relative transcriptional energy to mRNA. We also expect that the ribosomes should be actively translating those molecules given their short half-life. In support of our intuition, in both yeast and bacteria, we see the same strong correlation between the population’s average mRNA/rRNA ratio and the population’s growth rate that is expected given the growth law (**Figure S1a-i**; population growth rate explains 69% and 83% of variance in mRNA/rRNA ratio, respectively). But just as we observe for the absolute rRNA abundance, the correlation with growth rate becomes much weaker at the single-cell level (**Figure S1b-i**; population growth rate explains 9% and 5% of variation in mRNA/rRNA ratio respectively). Similar results are observed when looking at log-normalized and scaled scRNAseq counts (**Figure S1a, b-ii**) and ribosomal protein mRNAs (**Figure S1a, b-iii**). In sum, no matter how we normalize single-cell ribosome levels, we find large variation across cells that seems inconsistent with the idea that cells are optimizing their ribosomal content to support the maximum growth rate given their nutrient concentration.

This massive variation in single-cell ribosome levels also does not appear to be driven in large part by differences in cell size. For one, the normalization methods described above, such as taking the ratio of mRNA/rRNA and log normalizing, result in ribosome counts that are largely independent of cell size, and yet still vary across single cells (**Figure S1a, b**). Further, our analysis includes both haploid and diploid strains of yeast, which are known to differ in size by approximately 1.5X on average^53^. And yet, ploidy explains less than 1% of the variance in single-cell rRNA abundance in a categorical linear model (**Figure S2**). Finally, recent work in E. coli also demonstrates variation in ribosome concentration across single cells that is independent of cell size^37^. And recent work in bacteria suggests that smaller daughter cells tend to transiently have higher ribosome concentrations than their larger mothers^90^. In sum, this evidence suggests that single-cell ribosome levels can vary independently of cell size.

Finally, we found that variation in predicted cell cycle state^91^ also does not explain the variation we observe in the yeast data (**Figure S3**). This is consistent with other work suggesting that ribosome levels do not fluctuate with the cell cycle^7,11^.

### Estimating single-cell growth rates from transcriptional data

We inferred single-cell growth rates from gene expression profiles, or the genes and quantities of expression detected in each cell. Previous work reports linear relationships between gene expression and growth rate for the majority of *S. cerevisiae* genes^9^. Genes for which expression is higher during faster growth have positive slope values, and those for which expression is higher during slow growth have negative slopes. Using the same set of growth-predictive genes as in this previous study^9^, we used these slope values to weight normalized and scaled mRNA expression for each cell using the following formula with rRNA genes removed.

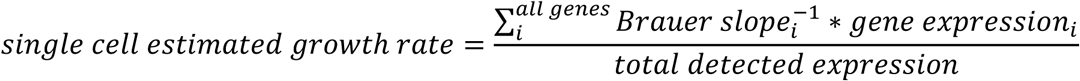

This equation produces an arbitrary, unitless number. The more positive the number, the more the transcriptome of that cell is indicative of growth-related expression and vice versa.

### Inferring variation in single cell growth rates (Figure 3a, b)

We used the inferred single-cell growth rates calculated by the above formula to ask if variation in single-cell growth rates positively correlates with the population growth rate, as does variation in single-cell ribosome levels. To do so, we took the coefficient of variation of the predicted single-cell growth rates for each population growth rate as a measure of growth rate heterogeneity. However, unlike the rRNA abundance heterogeneity, the heterogeneity in single-cell growth rate negatively correlates with population growth rate in yeast. This may suggest that variation in the growth of individual cells is not responsible for the spread of rRNA abundance we observe.

However, these measurements rely on growth correlation values estimated from bulk studies. Thus, we also performed a second analysis to confirm our results that does not rely on inference from scRNAseq data. For this analysis, we used an independent data set of single-cell *S. cerevisiae* growth measurements where the growth of thousands of yeast microcolonies was monitored over time using high-throughput microscopy^55^. We aggregated all microcolonies and averaged the growth rate for each timepoint to generate a proxy for population growth rate. Like our data, the mean growth rate generally decreases with time as nutrients are consumed. We then again calculated the heterogeneity using the coefficient of variation of the individual growth rates of the microcolonies as stratified by time point. Like our gene expression predicted growth rate data, the heterogeneity of single-cell growth rates declines as the population average increases. These negative correlations are the opposite of what we see in the rRNA data.

### “Apples to apples’’ rRNA quartile analyses (**Figure 4**)

We first calculated the minimum and maximum rRNA abundances for each timepoint and found the largest minimum and smallest maximum. The largest minimum and smallest maximum were used to create a bandwidth filter to allow for more direct comparisons of cells with similar rRNA amounts with representatives from each timepoint. Cells within this bandwidth were sorted by rRNA from least to greatest and quartiles were calculated. We then applied a Louvain clustering algorithm to these cells with a low enough resolution to only produce 2 to 3 clusters.

### Within-timepoint analyses including timepoints near steady state growth (Figure 5a)

Having observed more variation than expected in ribosome abundance, we further investigated to ask if these cell populations displayed heterogeneity in the rest of their transcriptomes. In yeast, we focused on the heterogeneity in the first and last timepoints. Theses samples correspond to the timepoints when the cells are at 9.2*10^6^ cells/mL and growing at an estimated 1.05 generations/hour and when cells are at 8.6*10^7^ cells/mL and growing at 0.09 generations/hour. In bacteria, we chose the timepoints when the cells were at OD600 of 1.0 and a growth rate of 1.24 generations/hour and OD600 of 1.3 and a growth rate of 1.14 generations/hour. This corresponds to the second and third timepoints in the replicate 1 bacterial series. These are the two timepoints in this replicate with the most recovered cells, significantly increasing the resolution of any subpopulations detected via clustering methods.

We then performed Louvain clustering as described above, and a differential expression analysis on these clusters and categorized the top 20 positive genes in each using SGD^84^, Metascape^92^, or SubtiWiki^85^. We used these categorizations to classify each cluster, and found they were dichotomized by growth and stress related genes. We sorted out the cells by barcode for each cluster and used density plots to visually compare the rRNA distributions.

### Identifying yeast cells expressing retrotransposons and bacterial cells expressing sporulation genes (Figure 5b, c)

When performing the differential expression analyses on the yeast timepoint 4 cells, we observed that the differentially expressed genes for cluster 0 included many transposable element genes for TY1 retrotransposons. These genes appeared to be co-transcribed in the cells with a less apparent stress response. This was unexpected, as retrotransposons are typically associated with stress, prompting us to examine transposable element gene expression across the yeast timepoints. We assembled a list of TE genes using SGD^84^, and calculated their combined sum expression for each cell in the normalized and scaled data. These values were mapped onto the UMAP plots using the FeaturePlot function in Seurat^83^.

When we studied the mRNA-rRNA ratio per cell, rather than rRNA counts, this revealed a major difference between bacterial replicate experiments. In replicate M14 from Kuchina and Brettner et al. 2021^36^, herein referred to as replicate 2, we observed a bimodal distribution of mRNA-rRNA ratios, where a subpopulation ranging from 16% to 50% of cells, depending on the timepoint, appeared to have a very high ratio of mRNA to rRNA (**Figure S7**). Upon performing a differential expression analysis, we observed that over 30% of the top 100 upregulated genes in these high mRNA-rRNA subpopulations were related to sporulation, leading us to believe these cells are in transition to a non-growing sporulation state. These putatively sporulating cells are present in every replicate 2 timepoint, and sample times in replicate 2 correspond to 6 timepoints ranging from OD600 0.5 to 3.2. We excluded this replicate from much of our analysis because we noticed that it appears to contain sporulating cells, however, the distribution of rRNA abundance follows a similar distribution to the other replicate (**Figure 3d**).

## ACKNOWLEDGEMENTS

Funding for this work was provided by a National Institutes of Health grant R35GM133674 (to KGS), an Alfred P Sloan Research Fellowship in Computational and Molecular Evolutionary Biology grant FG-2021-15705 (to KGS), and a National Science Foundation Biological Integration Institution grant 2119963 (to KGS). We would also like to thank Michael Lynch and Douglas Shepherd for helpful conversations, and the members of the ASU Biodesign Center for Mechanisms of Evolution and the Geiler-Samerotte Lab for their support and feedback.

## AUTHOR CONTRIBUTIONS

LB and KGS conceived of the study and contributed to writing the manuscript. LB performed the experiments and analyses.

