## Supplementary Materials (Figures) for "Single-cell heterogeneity in ribosome content and the consequences for the growth laws"

**SUPPLEMENTARY FIGURES**

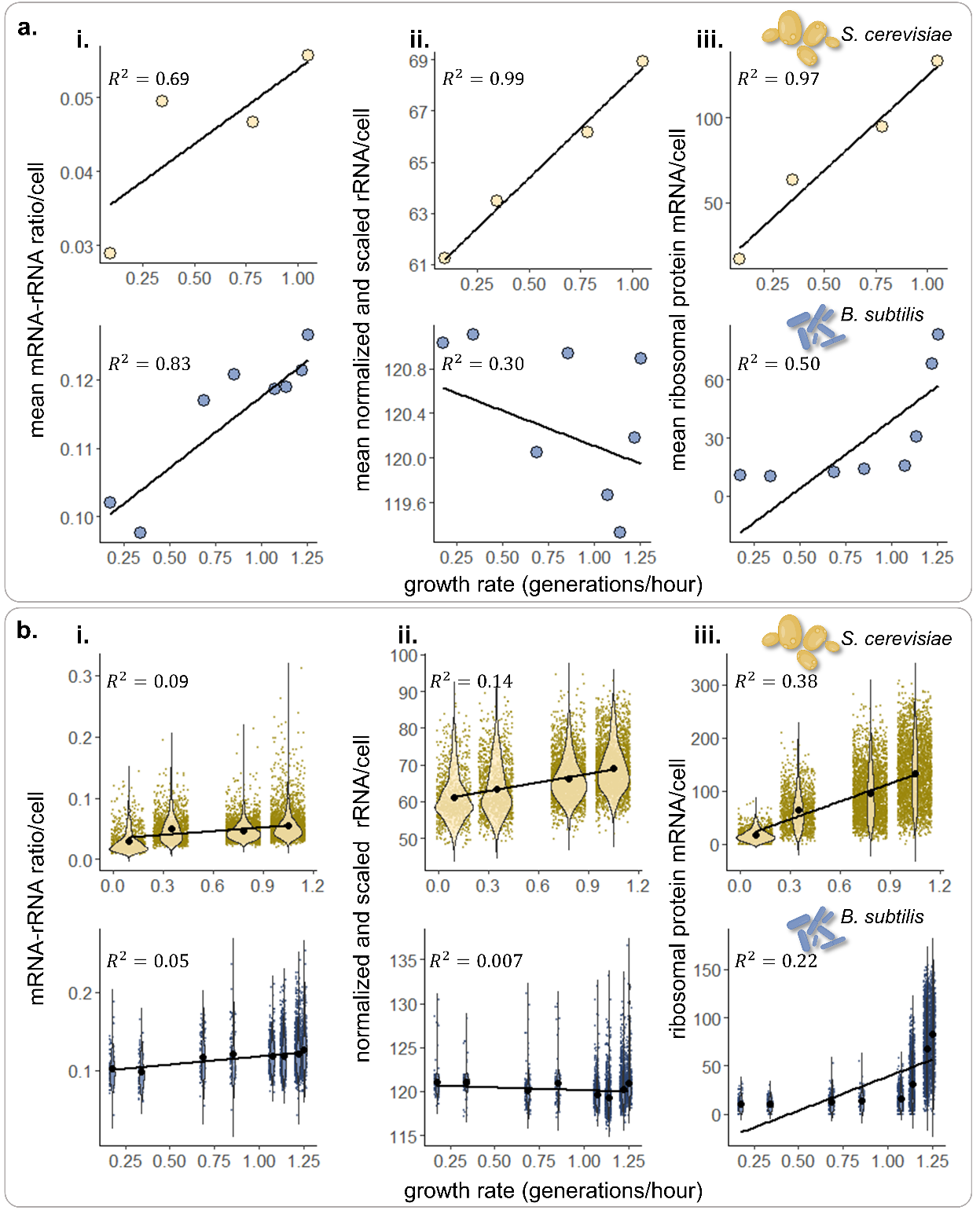

**Figure S1. Discordance in model predictability between means and distributions.** a.) i. Mean mRNA to rRNA ratio of QC processed but not normalized or scaled reads. ii. Mean normalized and scaled rRNA counts per cell as described in the Methods. iii. Mean ribosomal protein mRNAs per cell for normalized and scaled data with rRNA removed. b.) i-iii. The same analyses as in a.) except on the whole dataset distribution and not the means. R^2^s were derived from linear regression models of the variables displayed.

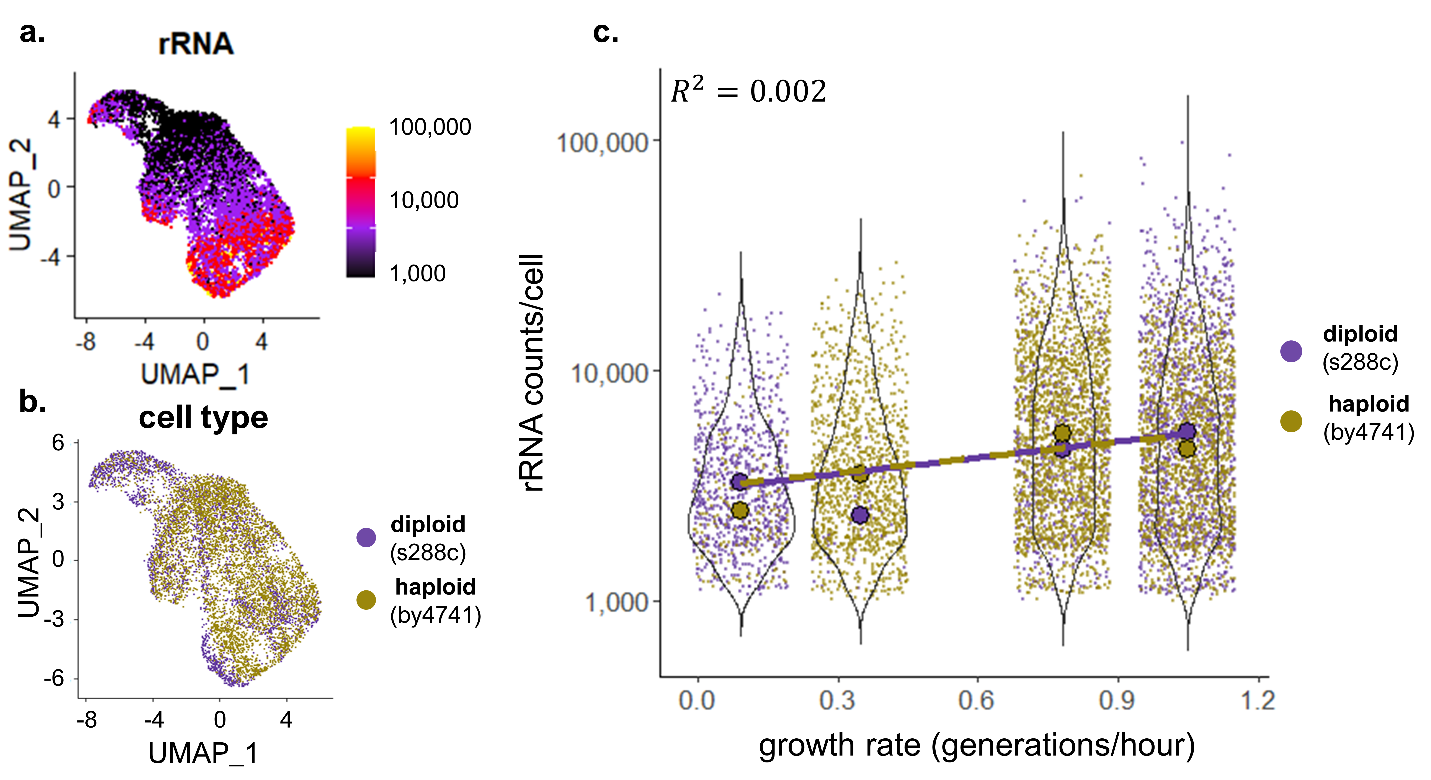

**Figure S2. Ploidy (as a proxy for cell size) is not predictive of rRNA in yeast.** a.) UMAP embedding of all cells from the yeast growth curve colored by rRNA abundance. b.) The same UMAP embedding colored by cell ploidy. c.) Means, trendlines, and distributions of rRNA/cell grouped by population growth rate and colored by ploidy. R^2^ was derived from a linear regression model with categorical predictors for ploidy.

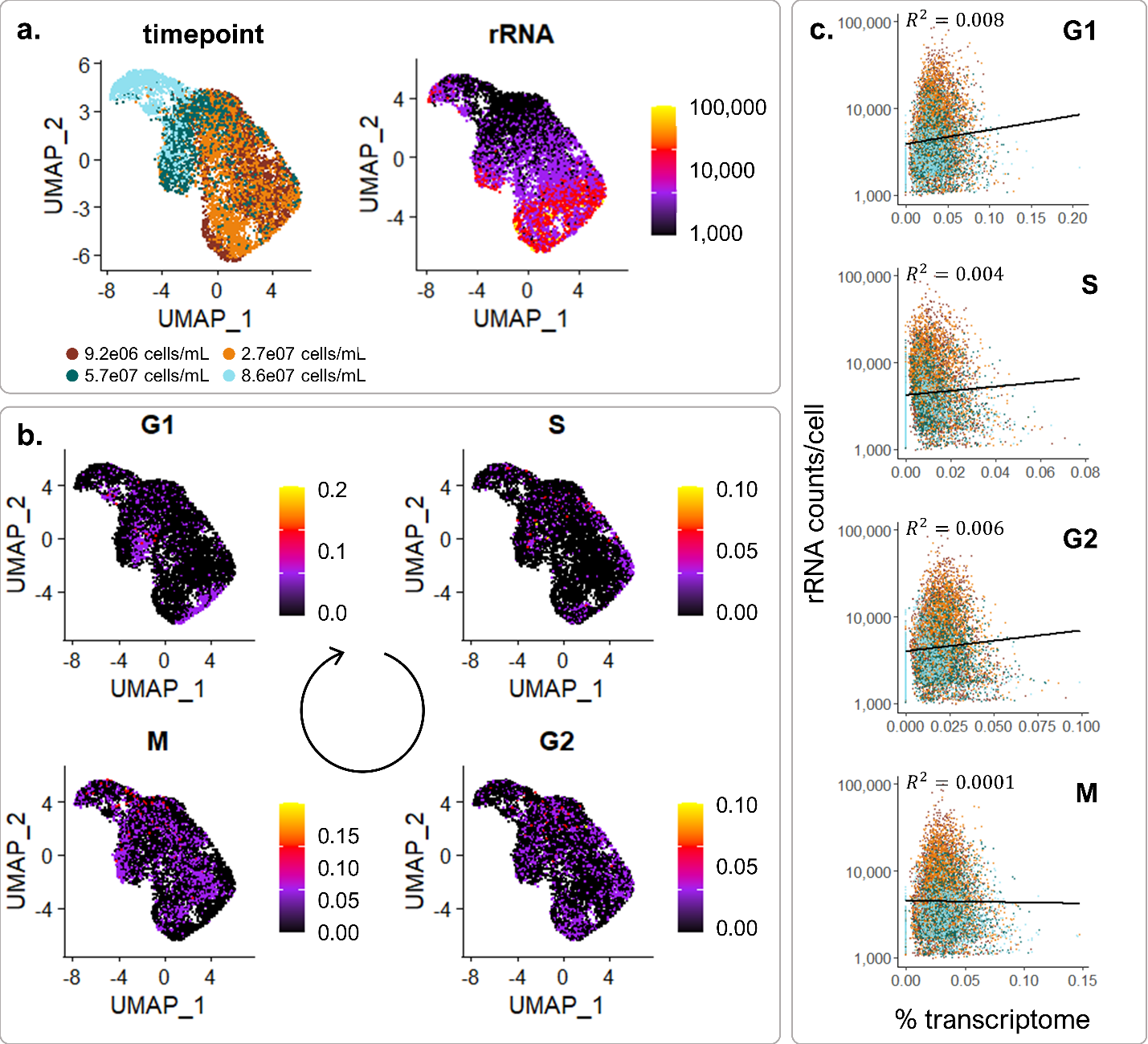

**Figure S3. Cell cycle state does not explain the variation we observe in rRNA abundance.** a.) UMAP embeddings of cells from all 4 timepoints in the yeast growth curve colored by sample point or rRNA counts. b.) UMAP embeddings colored by the percent of the mRNA transcriptome dedicated to cell cycle specific genes classified by Spellman et al. 1998^91^. c.) The percent of the transcriptome dedicated to cell cycle specific genes against the rRNA counts/cell. Datapoint colors represent the sample time to which each cell belongs as in panel a. R^2^s were derived from linear regression models of the variables displayed. An additional multivariable linear regression was calculated with the expression all the cell cycle genes against rRNA counts, and the resulting R^2^ = 0.017.

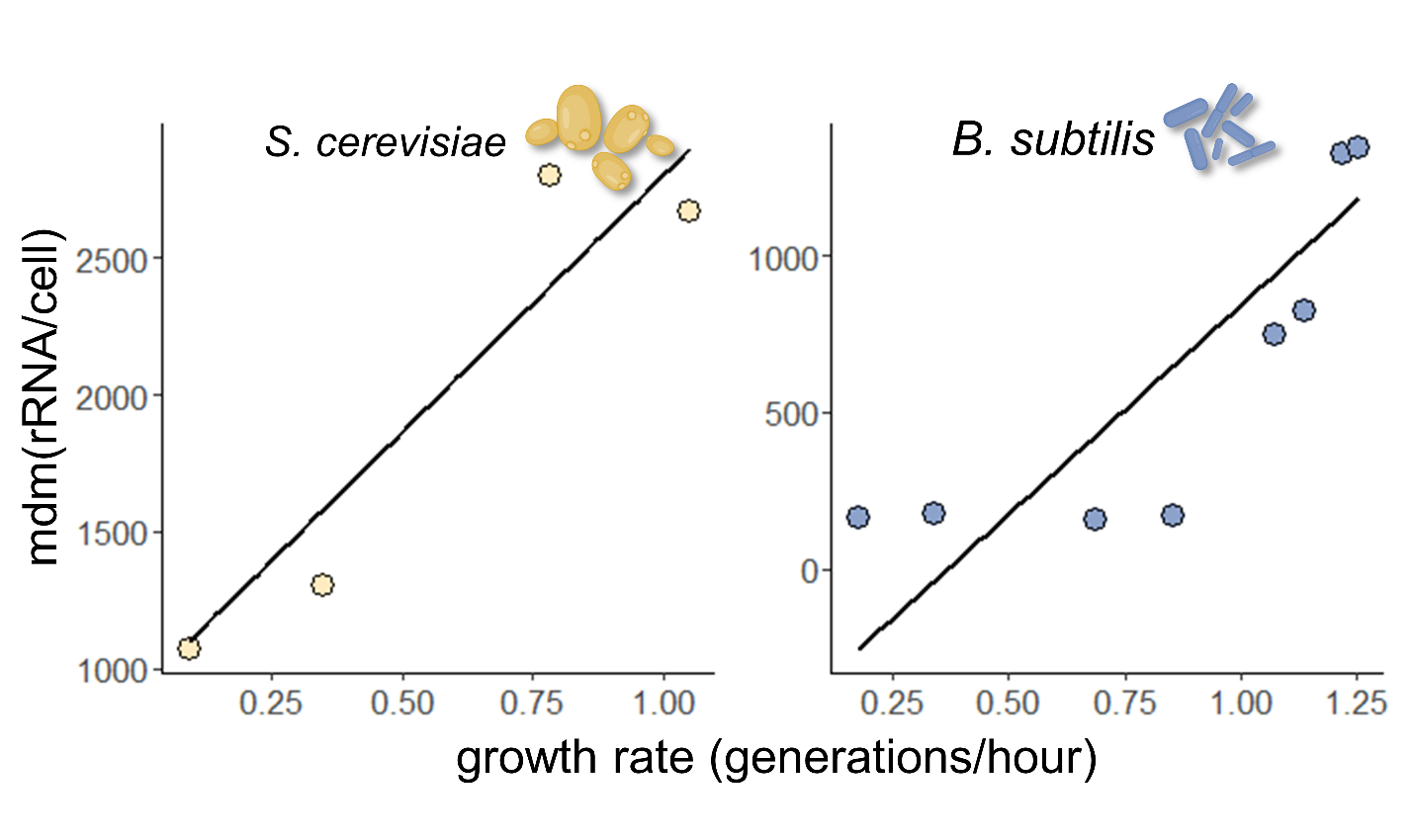

**Figure S4. Median deviation of the median rRNA per cell increases with population growth rate.** The mdm is an alternate measure of data spread to coefficient of variation for non-Gaussian distributions. The mdm was applied to rRNA counts for cells across each population growth rate for both yeast and bacteria. Trendlines are shown to indicate direction of change in mdm with population growth rate. This is the same trend we found when we used the coefficient of variation in **Figure 3**.

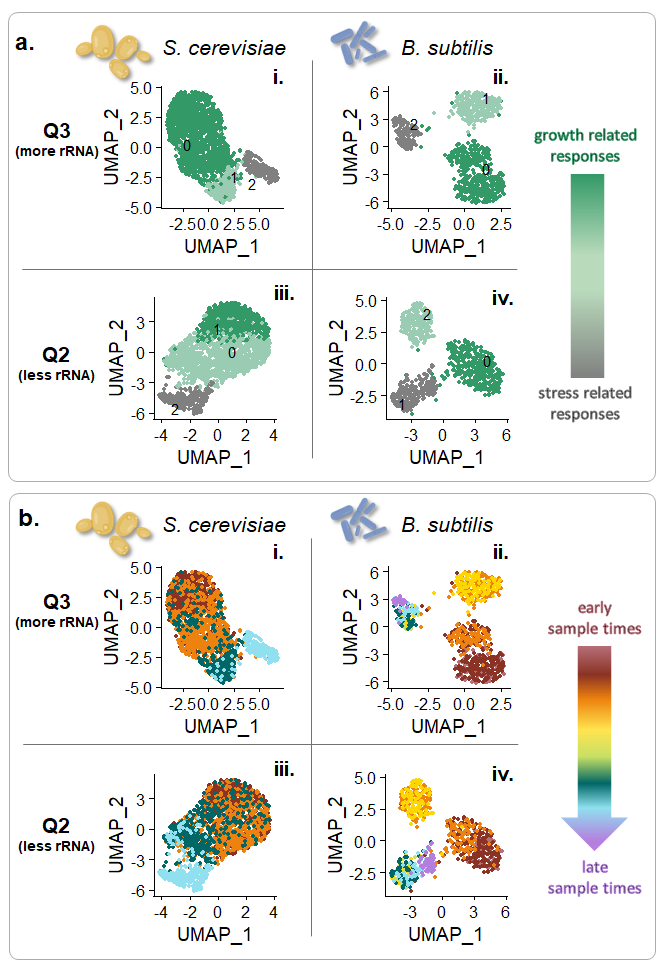

**Figure S5. Cells show differential subpopulations and signatures of sample timepoint in cells with intermediate ribosome levels a.)** UMAP embeddings of cells from the third (i-ii) and second (iii-iv) rRNA abundance quartiles colored by the Louvain cluster. **b.)** UMAP embeddings of cells from the third (i-ii) and second (iii-iv) rRNA abundance colored by sample timepoint.

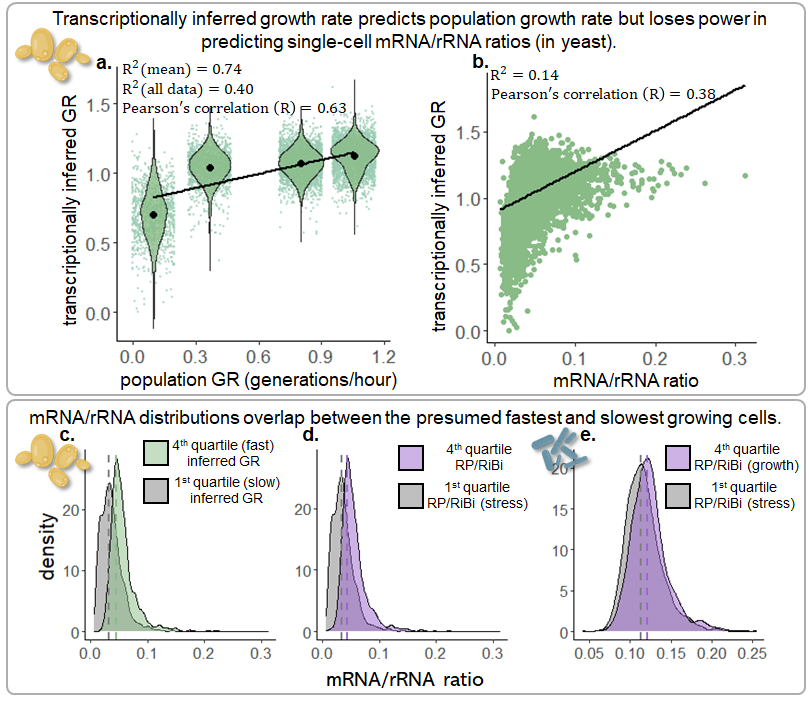

**Figure S6. mRNA/rRNA ratio in cells with growth associated transcriptional signatures does not widely differ from cells with stress signatures. a.)** Repeated from **Figure 2a.** Means and single-cell distributions of transcriptionally inferred growth rates against population growth rates (in yeast). **b.)** Transcriptionally inferred growth rate for each cell weakly correlates with corresponding mRNA/rRNA ratio. **c.)** mRNA/rRNA ratio density distributions for the cells with the 25% highest (green) and lowest (gray) inferred growth rates (in yeast) are largely overlapping. Dotted lines represent the density distribution maximum, or the largest mode. **d.)** and **e.)** Similarly overlapping distributions are observed for the cells with the highest percentage of the transcriptome dedicated to ribosomal proteins and ribosomal biogenesis genes (lavender) versus stress genes (gray) as defined in yeast^18^, and a similar cohort of genes for *B. subtilis* (Supplemental Table 1).

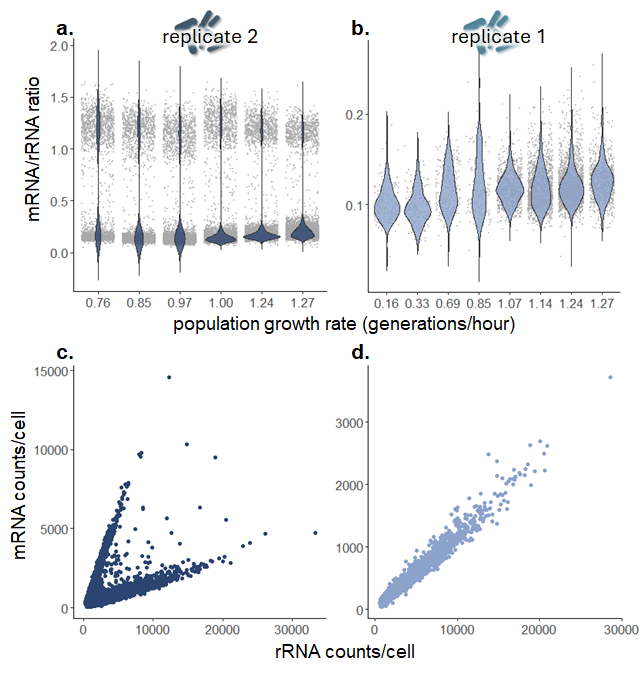

**Figure S7. Diverging of mRNA/rRNA across cells from two bacterial replicates. a and b.)** Cell distributions of mRNA/rRNA ratio across the growth curve for replicate 2 (a) and replicate 1 (b). Note the bimodal distributions appear only in replicate 2. **c and d.)** rRNA against mRNA for all bacterial cells and timepoints for replicate 2 (c) and replicate 1 (d). Note the cells with higher mRNA/rRNA ratio appear only in replicate 2.

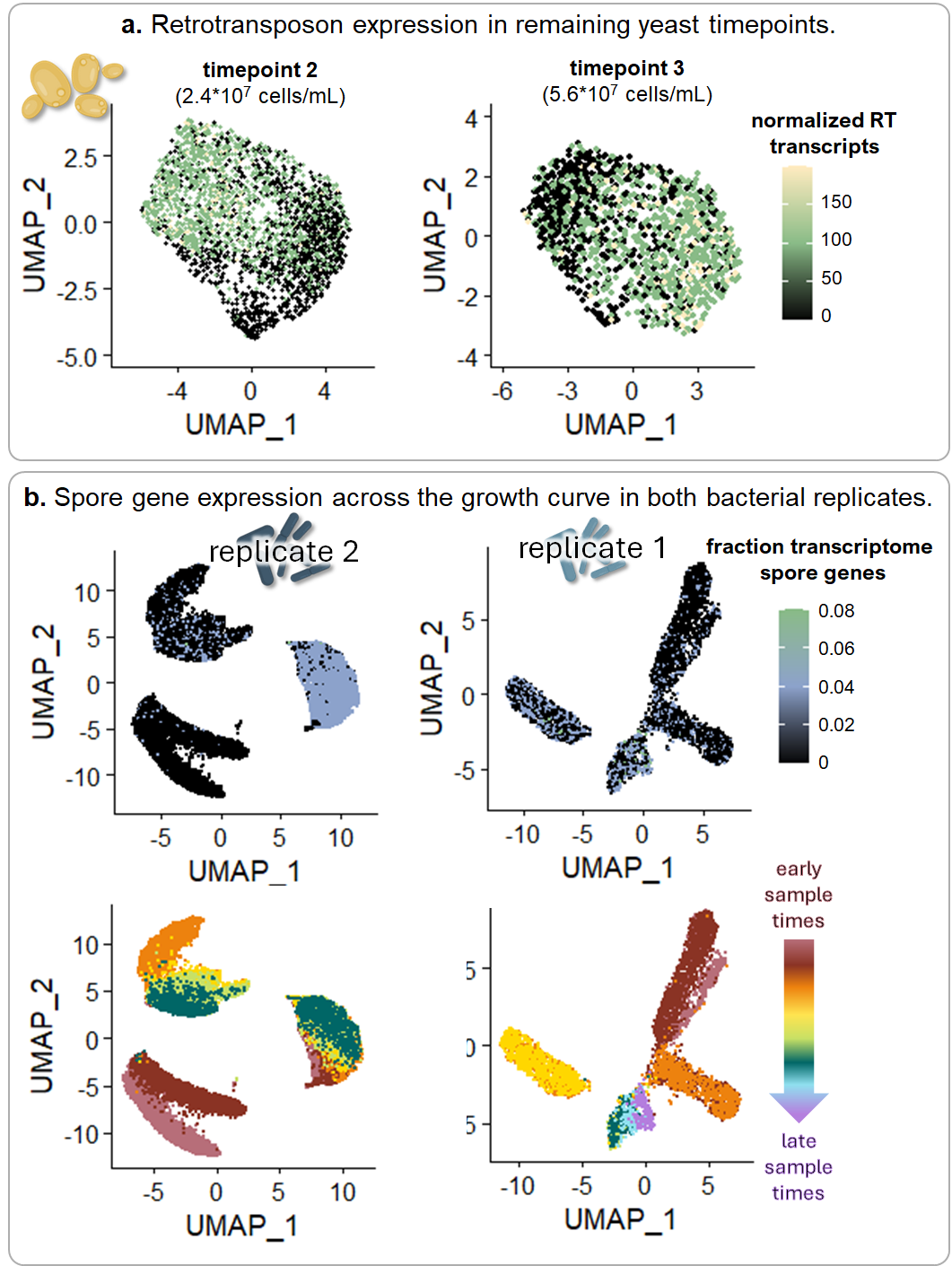

**Figure S8. Unexpected subpopulations are observed across the growth curve in both yeast and bacteria. a.)** UMAP embeddings of cells from timepoints 2 and 3 of the yeast growth curve colored by retrotransposon gene transcript abundance (Supplemental Table 1). **b.)** UMAP embeddings of bacterial cells from replicate 2 (left) and replicate 1 (right) colored by the fraction of the transcriptome dedicated to sporulation genes (Supplemental Table 1) (top) and by sample time (bottom).

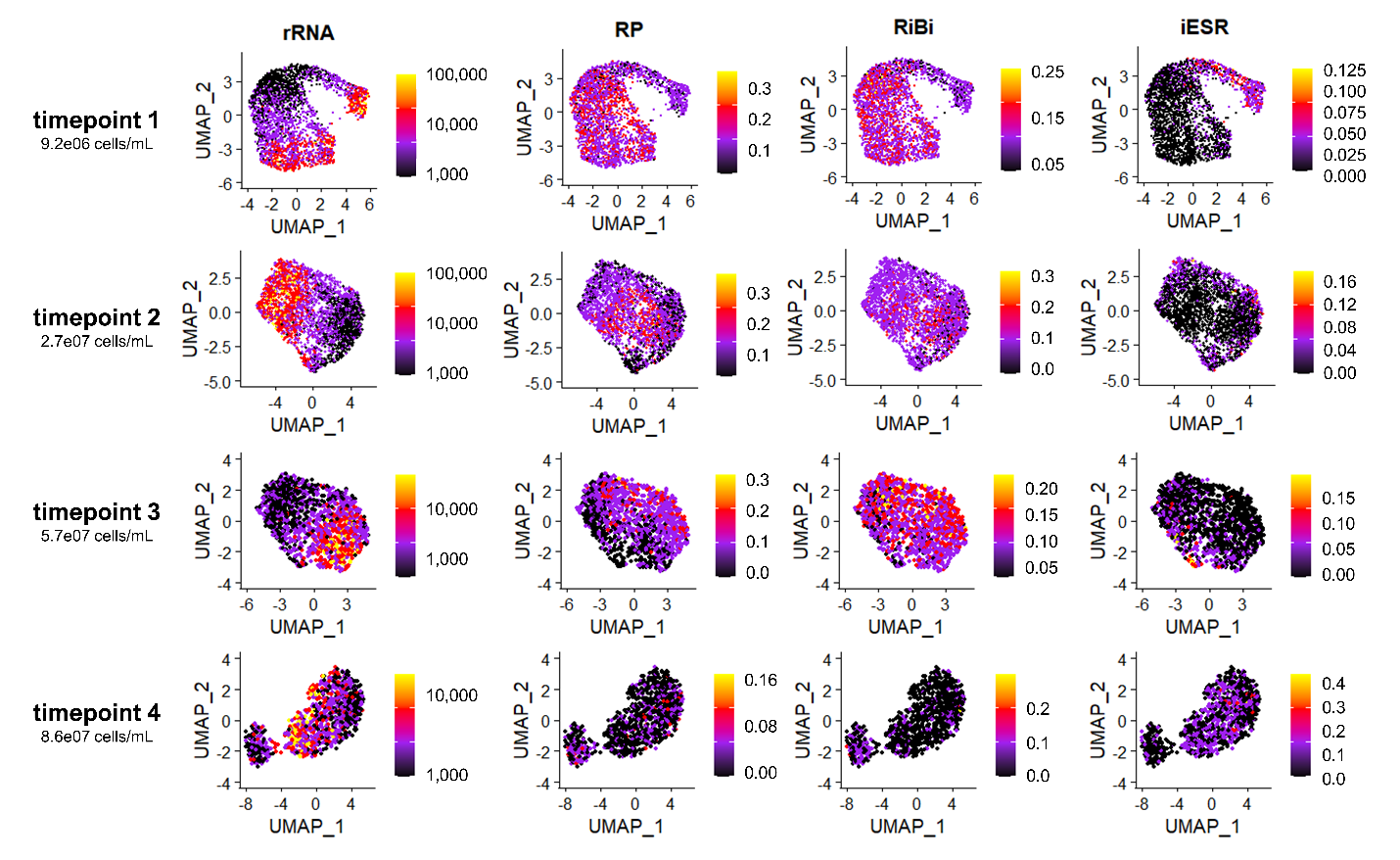

**Figure S9. Growth and stress response transcription across the yeast growth curve.** UMAP embeddings for each of the 4 timepoints from the yeast growth curve depicting rRNA counts, and percent of the mRNA transcriptome of growth response genes classified as ribosomal proteins (RP) and ribosomal biogenesis (RiBi), or the induced environmental stress response (iESR) as described in Gasch et al. 2000^18^. Note that cells expressing stress-related transcripts are present across all timepoints, as are cells expressing growth related transcripts.
